# Respiratory viruses induce ferroptosis-like features in lung epithelial cells

**DOI:** 10.1101/2025.10.29.685341

**Authors:** Ana L. Manzano-Covarrubias, Alessandra Tosato, Christina HT J. Mol-van der Veen, Philip Ahmadyar, Jai Walia, Jacqueline De Vries-Idema, Anke Huckriede, Amalia Dolga, Martina Schmidt, Karim Rafie

## Abstract

During infection, viruses can modulate various regulated cell death (RCD) mechanisms to evade host immunity and enhance replication. However, less is known about how viruses alter the recently discovered RCD ferroptosis, which is characterized by an iron-dependent accumulation of lipid peroxidation and mitochondrial fragmentation. Influenza A virus (IAV) H1N1 and human adenovirus type C5 (HAdV-C5) are two common causes of respiratory infections of the upper and lower respiratory tract and can lead to severe illness. While IAV has been shown to induce ferroptosis to support its replication, less is known whether this is also true for HAdV-C5. Here we perform a comparative study investigating ferroptosis features during IAV H1N1 and HAdV-C5 infections using alveolar (A549) and bronchial (BEAS-2B) epithelial cells. Our data reveal that HAdV-C5, similar to H1N1, induces lipid peroxidation in a time-dependent manner, which is partially suppressed in the presence of the ferroptosis inhibitor ferrostatin-1 (Fer-1). Strikingly, HAdV-C5 infections do not lead to changes in ferritin protein levels, which is responsible for iron storage. Furthermore, both H1N1 and HAdV-C5 infections trigger a profound mitochondrial network remodeling, comparable to that induced by the ferroptosis inducer RSL3, with effects varying by cell type and infection stage. These findings suggest that HAdV-C5, just like H1N1, can activate ferroptosis-like processes, highlighting a potential role for lipid peroxidation and mitochondrial alterations in adenoviral pathogenesis. Understanding this mechanism may uncover new therapeutic targets for viral respiratory infections.

## 1. Introduction

Regulated cell death (RCD) is an important host defense mechanism during viral infections restricting viral spreading and promoting viral clearance. It is also known that viruses can modulate RCD mechanisms to their benefit to promote replication and evade the immune system response [1–3]. Different types of RCD, including apoptosis, pyroptosis, and necroptosis have been extensively studied in the context of viral infection, however, much less is known about ferroptosis, an iron-dependent form of RCD [4–6]. Ferroptosis is characterized by several distinct hallmarks. First, the iron-dependent increase of lipid peroxidation, which results from a dysfunctional System Xc-(Xc-)/Glutathione Peroxidase 4 (GPX4) antioxidant pathway, ultimately reducing the levels of the antioxidant glutathione (GSH) [7]. Second, mitochondrial fragmentation, which is characterized by the loss of cristae as well as a smaller and more compact mitochondrial morphology [4, 7, 8]. Although, the role of mitochondria in ferroptosis is not well defined, a growing body of evidence indicates that they actively participate in ferroptosis by regulating mitochondrial reactive oxygen species (mtROS) production, iron and lipid metabolism, and redox balancing in the cell [9–11]. Further, recent studies reported that viral infections can disrupt these mitochondria-linked pathways to modulate ferroptosis [5, 6, 12, 13]. However, little information is available regarding whether virus-induced ferroptosis hallmarks are responsible for any changes in mitochondrial morphology.

Influenza A virus (IAV) is an RNA virus with a negative sense, single stranded, segmented genome that recognizes and binds to sialic acids widely present on the surface of nasal and bronchial epithelial cells [14]. IAV causes annual epidemics of acute respiratory infections and was the cause of four major pandemics [15]. Being one of the few viruses in which ferroptosis has been studied, evidence shows that influenza virus modulates the iron metabolism and GPX4 antioxidant axes to induce ferroptosis for its own benefit [16]. It was reported that swine influenza virus upregulates transferrin and transferrin receptor expression to promote intracellular iron accumulation and downregulates GPX4 expression, leading to lipid peroxide production and accumulation in alveolar epithelial A549 cells [17]. Similarly, H1N1 IAV increased ferrous and total intracellular iron and caused the mitochondria of infected cells to become smaller, with less cristae and compacted mitochondrial membranes in MLE-12 mouse lung epithelial cells [18]. Furthermore, H1N1 IAV uses hemagglutinin to form ferritin-nuclear receptor coactivator 4 (NCOA4) condensates, promoting ferritinophagy to increase free intracellular iron [19] and uses transferrin receptors to enter the host to enhance its replication [20].

Human adenoviruses (HAdV) are a large group of non-enveloped, double-stranded DNA viruses most of which use the coxsackievirus and adenovirus receptor (CAR) protein present in epithelial tight junctions for viral attachment and entry [21, 22]. With over 110 types classified into seven species (A-G), HAdVs are widely spread, and are known to cause severe disease in immunocompromised patients [23, 24]. HAdV regulation of apoptosis, necrosis, and autophagy have been widely studied [25], but whether ferroptosis is a common RCD mechanism across HAdVs remains unclear. HAdV species C (HAdV-C) account for up to 10% of pediatric and up to 7% of adult respiratory infections and have been known to cause epidemics in congregate living spaces [26, 27]. HAdV-C5, is among the most common HAdVs associated with human disease, especially in immunocompromised patients [28], but whether or not it also causes ferroptosis-like features has not yet been reported. The adenovirus early region 1A gene has been reported to inhibit the expression of ferritin and sensitizing infected cells to oxidative stress [29]. Recent research on fowl adenovirus serotype 4, a virus that causes hydropericardium syndrome and liver damage in poultry, reported on the induction of ferroptosis features including elevated levels of malondialdehyde (MDA), an end product of lipid peroxidation, as well as reduced levels of GSH, reduced GPX4 activity, and a reduced expression of ferritin heavy chain (FTH1) in a chicken hepatocellular carcinoma cell line (LMH) [30]. Similar findings have been reported for HAdV 36, which caused an increase of MDA in liver tissue of embryonated chicken eggs [31] and a decrease in mitochondrial mass in a dose-dependent manner in infected human skeletal muscle cells [32].

In general, as mentioned above, research with focus on the interaction between ferroptosis and virus infections is limited. Therefore, we investigated in this study whether HAdV-C5 infections induce ferroptosis-like features in human bronchial (BEAS-2B) and alveolar (A549) lung epithelial cell lines. As IAV are among the few viruses in which ferroptosis signaling mechanisms have been studied [6, 16, 17, 19, 20, 33], we used the H1N1 influenza virus as the model virus to determine whether HAdV-C5 is capable of activating this form of RCD. Our findings indicate that HAdV-C5 causes an increase in lipid peroxidation seemingly leaving the expression of iron storage protein ferritin unaffected. Importantly, we demonstrate changes in the mitochondrial network in lung epithelial cells caused by both H1N1 and HAdV-C5 infection.

## 2. Materials and methods

### 2.1 Cell culture and treatment

Adenocarcinomic human alveolar epithelial (A549) cells (ATCC CCL-185) and human bronchial epithelial (BEAS-2B) cells (ATCC CRL-3588) were cultured in Roswell Park Memorial Institute (RPMI)-1640 medium with GlutaMAX^TM^ Supplement (RPMI 1640 Medium, GlutaMAX^TM^ Supplement; #61870044; Gibco, Thermo Fisher Scientific, Netherlands) supplemented with 10% (v/v) heat-inactivated fetal bovine serum (HI-FBS, Hyclone, USA) and 2% (v/v) of antibiotics (Penicillin-Streptomycin 5000 U/mL, #11528876, Gibco, Thermo Fisher Scientific, Netherlands) and kept in a humidified incubator at 37 °C and 5% (v/v) CO_2_. Cells were detached with trypsin-EDTA (Cat#L0930, Biowest) and seeded in appropriate cell culture plates. During treatment cells were maintained in starvation media (RPMI-1640 medium with GlutaMAX^TM^ Supplement with 1% (v/v) HI-FBS and 2% (v/v) antibiotics) [34, 35]. RAS-selective lethal small molecule 3(RSL3, #S8155, Selleckchem, USA) at a concentration of 10 μM for A549 and 250 nM for BEAS-2B was used to trigger ferroptosis, unless otherwise indicated. Ferrostatin-1 (Fer-1; #SML0583, Sigma-Aldrich) was used at 5 μM as a ferroptosis inhibitor [19, 36].

### 2.2 Viruses and infection

Influenza virus strain A/Puerto Rico/8/34 (H1N1/PR8) was kindly provided by Prof. Dr. Anke Huckriede (Department of Medical Microbiology, University of Groningen. Netherlands). The virus was propagated in embryonated chicken eggs, purified by sucrose density gradient centrifugation, and titrated on MDCK cells as described [37]. Human adenovirus type 5 (HAdV-C5) was purchased from The Gene Vector Core at the Baylor College of Medicine. Virus stocks were prepared by amplification in A549 cells followed by titre determination by plaque assay in A549 cells [38].

At 80% confluence, cells were infected with H1N1 or HAdV-C5 at multiplicity of infection of 1.0 (MOI 1) in infection medium (RPMI-1640 medium with GlutaMAX^TM^ Supplement with 2% (v/v) antibiotics) and incubated for one hour with H1N1 and two hours with HAdV-C5 to allow the adsorption of the virus, then cells were washed three times with phosphate buffered saline (PBS) and starvation medium was added. Infected cells were incubated for the desired time in a humidified incubator at 37 °C and 5% (v/v) CO_2_.

### 2.3 Immunofluorescence staining of viral proteins

A549 and BEAS-2B cells were seeded at 20,000 cells/well onto 48-well plates and infected with influenza virus (H1N1/PR8) virus as described above. After 0, 4, 6, 8, 18 and 24 hours post infection (hpi) cells were washed with PBS and fixed with 4% paraformaldehyde (PFA) solution at room temperature (RT) for 15 min. After washing three more times with PBS, cells were incubated overnight at 4 °C in blocking buffer (0.1% (w/v) saponine, 1% (w/v) bovine serum albumin in PBS). On the next day, cells were stained with primary antibody against influenza A nucleoprotein (NP) (Cat. #MAB8257, Millipore, 1:750 in blocking buffer) for 1 h at RT. Following incubation cells were washed three times with blocking buffer and then incubated with Alexa Fluor 568-conjugated donkey anti-mouse IgG (H+L) secondary antibody (Cat. #A10037, Invitrogen) for 45 min at RT [39].

A549 and BEAS-2B cells were seeded at 20,000 cells/well onto 48-well plates and infected with HAdV-C5 virus as described above. After 0, 24, 48, 72, and 96 hpi cells were washed with PBS, fixed with ice-cold methanol, and incubated at -20 °C for 10 min. Methanol was removed and primary antibody against adenovirus hexon protein (Cat. #MAB8052, Millipore, 1:500 in PBS) was added and incubated for 45 min at RT. After incubation cells were washed three times with PBS and incubated with Alexa Fluor 568-conjugated donkey anti-mouse IgG (H+L) secondary antibody (Cat. #A10037, Invitrogen) for 30 min at RT [40].

After incubation with secondary antibody, cells were washed three times with PBS and nuclei were stained with DAPI (Cat. #ab104139, Abcam). Fluorescence was detected on a Nikon Inverted Research Fluorescence Microscope ECLIPSE Ti2-E at constant exposure of 100 ms. At least 5 images of each well were captured at a nominal magnification of 20x. H1N1 and HAdV-C5 infected cells were quantified by counting nucleoprotein positive (NP+) cells and hexon protein positive (Hexon+) cells, respectively, and DAPI positive (DAPI+) cells using ImageJ [41]. Results are shown as percentage of NP+/DAPI+ or Hexon+/DAPI+ cells.

### 2.4 Cell morphology

After the desired time post infection or 24 hour treatment with RSL3, cells were washed with PBS and fixed with 4% PFA solution at RT for 15 min. Cells were washed three more times with PBS and imaged using a Nikon Eclipse Ti-S microscope at a nominal magnification of 10x.

### 2.5 Cell metabolic activity assay

Cell metabolic activity was measured using the 3-(4,5-Dimethyl-2-thiazolyl)-2,5-diphenyl-2H-tetrazolium bromide (MTT) assay (#M5655, Sigma-Aldrich, Netherlands). A549 cells were seeded at 9,000 cells/well and BEAS-2B cells at 10,000 cells/well onto a 96-well plate. After treatment time, MTT solution was added at a final concentration of 0.5 mg/mL and incubated for 1 h at 37 °C and 5% (v/v) CO_2_. The MTT solution was completely removed and to dissolve the formed formazan 70 µL dimethyl sulfoxide (DMSO) was added and incubated for 45 min at 37 °C at a constant shaking rate of 120 rpm. Lastly, absorbance was measured at wavelengths 570 and 630 nm using a Synergy^TM^ H1 Multi-Mode Reader (BioTek, USA) [42].

### 2.6 Cell death assay

Cell death was determined by staining the cells with propidium iodide (PI) and Hoechst 33258. A549 and BEAS-2B cells were seeded at 20,000 cells/well onto 48-well plates and infected with influenza virus (H1N1/PR8), HAdV-C5 virus, or treated with RSL3 as described above. At the indicated times post infection, PI (PI; #V13242; Fisher Scientific, Landsmeer, the Netherlands) and Hoechst 33258 (#H1398; Invitrogen, the Netherlands) were added to each well at a final concentration of 1 µM and 1 µg/mL, respectively. Cells were incubated at 37 °C and 5% (v/v) CO_2_ for 10 min and then imaged using a Nikon Inverted Research Fluorescence Microscope ECLIPSE Ti2-E at a nominal magnification of 20x. The number of PI+ and Hoechst+ cells was quantified using Fiji [43]. The average number of Hoechst^+^ cells in the control group was calculated 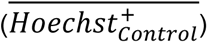, then for each image the percentage of cell death relative to control was calculated the following way:

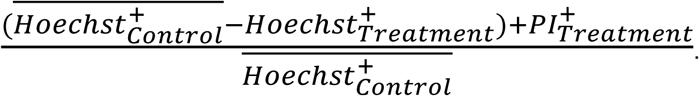

Ten images per condition were analyzed.

### 2.7 Lipid peroxidation

Lipid peroxidation was determined using BODIPY 581/591 C11 Lipid Peroxidation Sensor ((#D3861, Fisher Scientific, Landsmeer, the Netherlands) [42, 44, 45]. A549 and BEAS-2B cells were seeded at 20,000 cells/well onto 48-well plates and infected with IAV (H1N1/PR8) or HAdV-C5 or treated with RSL3 as described above. At the indicated times post infection, BODIPY 581/591 C11 and Hoechst 33258 dye were added to each well at a final concentration of 1.5 µM and 1 µg/mL, respectively. Cells were incubated at 37 °C and 5% (v/v) CO_2_ for 15 min and then imaged using a Nikon Inverted Research Fluorescence Microscope ECLIPSE Ti2-E at a nominal magnification of 20x. Ten images per condition per biological replicate were taken and analyzed using Fiji [43].

### 2.8 Malondialdehyde assay

Malondialdehyde (MDA) was measured as a marker of lipid peroxidation by the thiobarbituric acid reactive substances (TBARS) assay [42, 46]. A549 cells were seeded at 275,000 cells/well and BEAS-2B cells at 320,000 cells/well onto a 6-well plate. Infection was done as described above. Fer-1 treatment was added one hour prior to infection and removed at 0 hpi. RSL3 treatment was added at the same time of infection. At the indicated times post-infection cells were washed three times with PBS and collected by scraping in ice-cold water. Cells were lysed by sonication and freeze-thaw cycles for a total of three times. 50 μL of cell lysate was mixed with 100 μL ice-cold 10% trichloroacetic acid (TCA, Cat.#1.00807, Merck) and incubated for 15 min on ice. Samples were centrifuged for 15 min at 5,000x g at 4 °C and 100 µL of supernatant was mixed with an equal volume of 0.75% 2-thiobarbituric acid (TBA, Cat. #T550, Merck). An MDA (1,1,3,3-tetramethoxypropane, Cat.#108383, Merck) standard curve (0-500 μM) was prepared the same way. Samples and standards were immediately boiled at 95 °C for 15 min then allowed to cool to RT. Fluorescence was measured at λ_Ex_/λ_Em_ = 533/559 nm using a BioRad CFX96 Real-Time PCR System. Total protein concentration of samples was measured using the Pierce BCA Protein Assay Kit (Cat. #23335, Thermo Scientific). MDA concentrations in samples were calculated from the MDA standard curve and normalized by their respective total protein concentration.

### 2.8 Western blotting

A549 cells were seeded at 275,000 cells/well and BEAS-2B cells at 320,000 cells/well onto a 6-well plate. Infection was done as described above. Samples were collected at the indicated times post-infection in lysis buffer (0.25 M D-Manitol, 0.05 M Tris-Base, 1 mM EDTA, 1mM EGTA, pH 7.8 and a mixture of protease inhibitors: 1 mM Na3VO4, 1 mM NaF, 10 μg/mL leupetin, 10 μg/mL pepstatin A, 10 μg/mL apoprotein and 1 mM DTT). Total protein was quantified by Pierce BCA Protein Assay Kit (Cat. #23335, Thermo Scientific). Equal amounts of protein were loaded and separated by 12% SDS polyacrylamide gel electrophoresis, transferred onto nitrocellulose membranes, and blocked with 5% milk. Subsequently, membranes were incubated overnight at 4 °C with primary antibodies against ferritin light chain (FTL) (1:1000, Cat. #ab69090, Abcam) and GAPDH (1:3000, Cat. # sc-47724, Santa Cruz Technology) in 1% milk. Membranes were washed thoroughly and incubated with horse-radish peroxidase-conjugated secondary antibodies (anti-rabbit IgG, 1:3000; anti-mouse IgG, 1:3000, respectively) at RT for 1 h. Antibody-labeled proteins were detected using a Western Lightning ECL-plus Chemiluminescent Substrate detection kit (Cat. #50-904-9326, FisherScientific). ImageJ [41] was used for the densitometric analysis of the bands. Expression of ferritin was normalized to GAPDH.

### 2.9 Mitochondrial staining and morphology analysis

A549 and BEAS-2B cells were seeded at 40,000 cells/well onto glass coverslips in a 24-well plate followed by the virus infections as outlined above. Mitochondria were labeled by incubating the cells with 100 nM MitoTracker^TM^ Red FM (Invitrogen, Oregon City, OR, USA, #M22425) for 30 min. Following incubation time, cells were washed three times with PBS and fixed with 4% PFA solution at RT for 15 min. After washing three more times with PBS, coverslips were mounted on glass slides using mounting media with DAPI (Cat. #ab104139, Abcam). At least ten images per condition were acquired using a Nikon Inverted Research Fluorescence Microscope ECLIPSE Ti2-E at a nominal magnification of 60x and analyzed using ImageJ software.

Mitochondrial footprint was analyzed using the *MiNA* plugin (https://github.com/StuartLab/MiNA/blob/master/README.md) for Fiji [43, 47]. At least ten single cell images were analyzed for each condition. Branch length per mitochondrion, branches per mitochondrion, branch junctions per mitochondrion and mean form factor were measured from binarized images using the *Mitochondria Analyzer* plugin (https://github.com/AhsenChaudhry/Mitochondria-Analyzer) [43, 48]. Optimal threshold parameters were first determined and maintained for subsequent analysis for all images.

### 2.10 Statistical analysis

GraphPad Prism software (version 8.4.2, GraphPad Software Inc., La Jolla, CA, USA) was used to create all graphs and analyze data. All data are expressed as mean ± standard deviation (SD) for all experiments repeated as indicated in the figure legends. Analysis was performed using one-way ANOVA followed by Dunnett’s multiple comparisons or by Tukey’s post hoc multiple comparisons test. Significance is indicated by: *p < 0.05, **p < 0.01, ***p < 0.001, ****p < 0.0001, or otherwise not significant (n.s.).

## 3. Results

### 3.1 H1N1-induced cell death of epithelial cells leads to increases in lipid peroxidation and ferritin expression

It was previously demonstrated that influenza virus induces ferroptosis in A549 cells [17, 19]. We evaluated whether the H1N1/PR8 strain causes ferroptosis in both alveolar (A549) and bronchial (BEAS-2B) epithelial cells. We first assessed infection dynamics by infecting cells with H1N1/PR8 at a multiplicity of infection (MOI) of 1 and monitored nucleoprotein (NP) expression at 0, 4, 6, 8, 18 and 24 hours post infection (hpi) by immunostaining (supplementary Fig. 1). In A549 cells, NP-positive (NP+) cells reached 84.7% by 18 hpi and 93.5% by 24 hpi (Fig. 1A and 1B). In contrast, BEAS-2B cells showed faster NP production, with 58.9% cells already being NP+ at 4 hpi and peak infection levels (>90%) by 6 hpi (Fig. 1C and 1D).

**Fig. 1.**
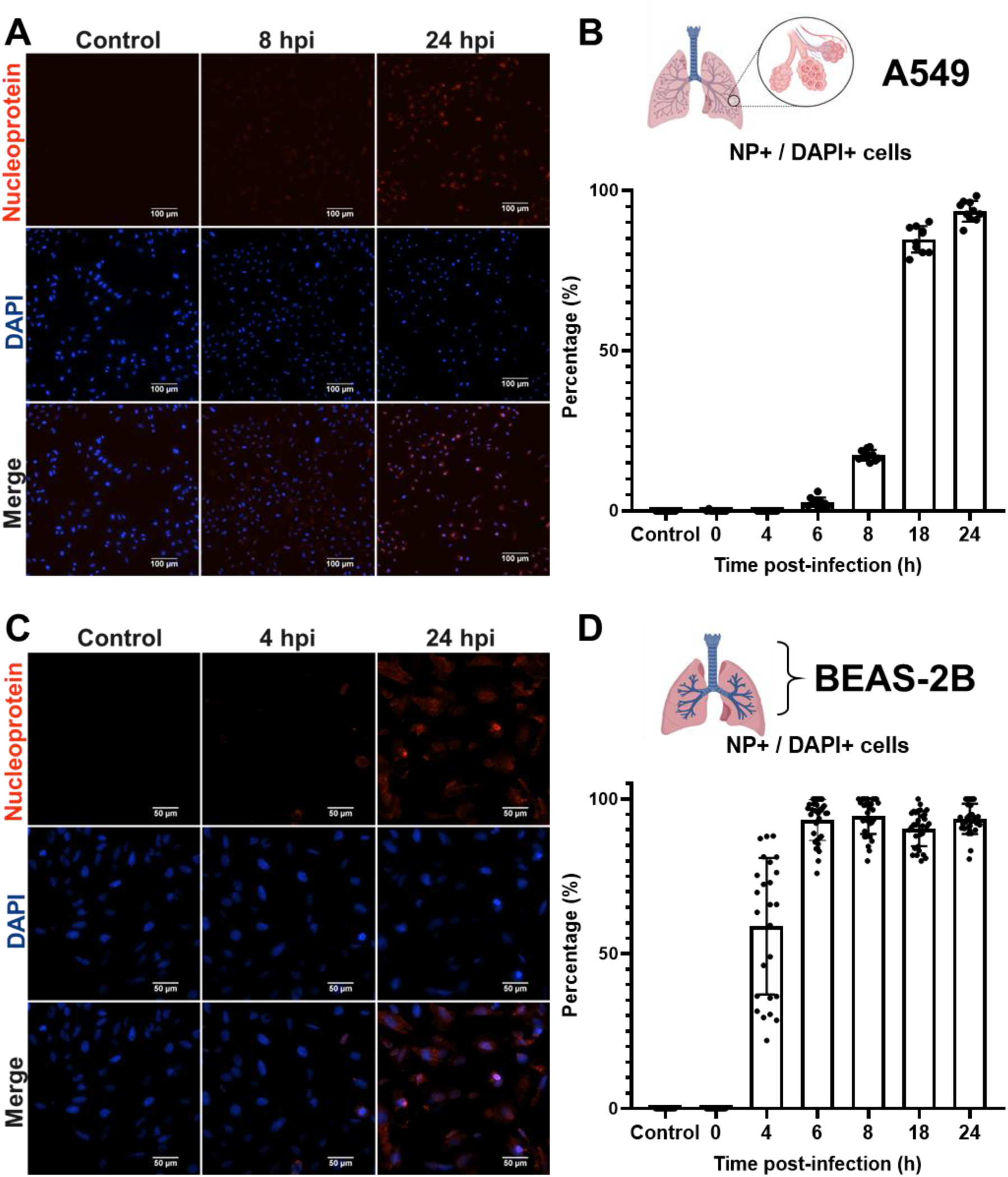
H1N1 replication in lung epithelial cells. Fluorescence images of (A) A549 and (C) BEAS-2B cells infected with H1N1/PR8 influenza virus at a MOI = 1, fixed and stained at the indicated time post infection taken at a nominal magnification of 20x. Staining was performed with an anti-influenza nucleoprotein (NP) antibody and counterstaining with DAPI. Red: NP positive cells; Blue: nucleus (scale bar = 100 µm for A549 cells and 50 µm for BEAS-2B cells). Quantitative analysis of NP positive and DAPI positive (B) A549 and (D) BEAS-2B cells was done on at least 5 images per condition per biological replicate using ImageJ. Results are shown as percentage of NP^+^/DAPI^+^ cells (mean ± SD).

Since both cell lines showed maximal infection by 24 hpi, we continued to use this time point for subsequent experiments, with RSL3, a known ferroptosis inducer [49, 50], used as a positive control. In conjunction with increased NP production, morphological changes and a reduction in cell number were observed in both H1N1-infected and RSL3-treated cultures for both A549 (Fig. 2A) and BEAS-2B (Fig. 2E) cells. To investigate whether virus infection leads to changes in the metabolic activity, we performed MTT assays, which showed a significant reduction in cell metabolic activity in H1N1-infected A549 (73.48% ± 6.83%) (Fig. 2B) and BEAS-2B (70.03% ± 1.62%) cells (Fig. 2F) compared to their respective controls. Given that the MTT assay measures cell metabolic activity and only indirectly provides insight into cell viability, we sought to quantified cell death by using propidium iodide (PI) and Hoechst 33258 staining. These results confirmed significant cell death in both A549 (50.1% ± 8.19%) (Fig. 2C and 2D) and BEAS-2B (49.1% ± 2.94%) (Fig. 2G and 2H) cells following H1N1 infection, which correlates with our findings in the MTT assay. For comparison, RSL3 treatment induced cell death levels of 56.95% ± 4.21% in A549 and 66.98% ± 3.32% in BEAS-2B cells.

**Fig. 2.**
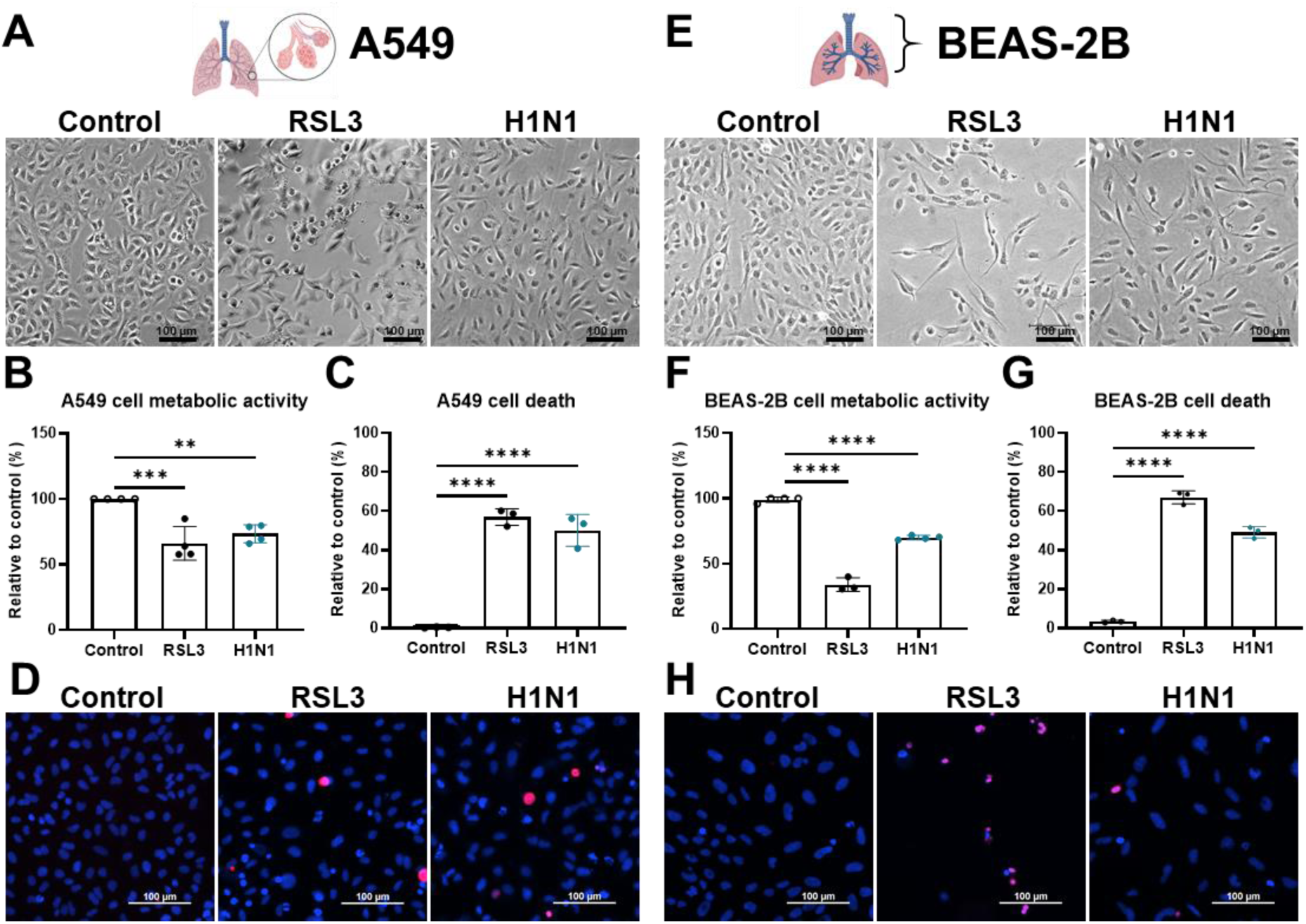
H1N1 causes cell death in lung epithelial cells. Representative brightfield microscope images of (A) A549 and (E) BEAS-2B cells infected with H1N1/PR8 influenza virus at a MOI = 1 at 24 hpi or treated with RSL3 (A549: 10 µM; BEAS-2B: 250 nM) (scale bar = 100 µm). MTT assay as measurement of cell metabolic activity of viable cells (B) A549 and (F) BEAS-2B cells. Cell metabolic activity of the control group was defined as 100% activity. Each dot represents a biological replicate with at least 6 technical replicates per condition. Results are shown as percentage relative to the control (mean ± SD). Cell death of (C) A549 and (G) BEAS-2B quantification by PI and Hoechst double staining and representative images (D: A549 and H: BEAS-2B) taken at a nominal magnification of 20x. The number of dead cells was quantified according to the equation stated in the methods section. In short, the total number of dead cells is calculated as the sum of PI+ cells and the difference between the average number of DAPI+ cells in the control and the number of DAPI+ cells in the treatment, divided by the average number of DAPI+ cells in the control. Red: PI positive, dead cells; Blue: nucleus (Scale bar = 100 µm). Each dot represents the average of ten images per condition. Results are shown as percentage of dead cells relative to number of nuclei present in the control group (mean ± SD). Statistical analysis was performed using one-way ANOVA followed by Dunnett’s multiple comparisons test. **p < 0.01, ***p < 0.001, ****p < 0.0001. Results are of at least three biological replicates.

A central feature of ferroptosis is the excessive and uncontrolled peroxidation of polyunsaturated fatty acids (PUFAs) present in cell membranes, a process that precedes and contributes to cell death [51, 52]. To assess this in our model, we measured lipid peroxidation in H1N1-infected cells at 24 hpi by staining A549 (Fig. 3A) and BEAS-2B (Fig. 3B) cells with the fluorescent probe BODIPY^TM^ 581/591 C11. Oxidation of the polyunsaturated butadienyl portion of the dye causes a fluorescence emission shift from red (∼590 nm) to green (∼510), allowing for the quantification of oxidized lipids to non-oxidized lipids. The ratio of green to red fluorescence intensity then allows for the quantification of lipid peroxidation (Fig. 3C).

**Fig. 3.**
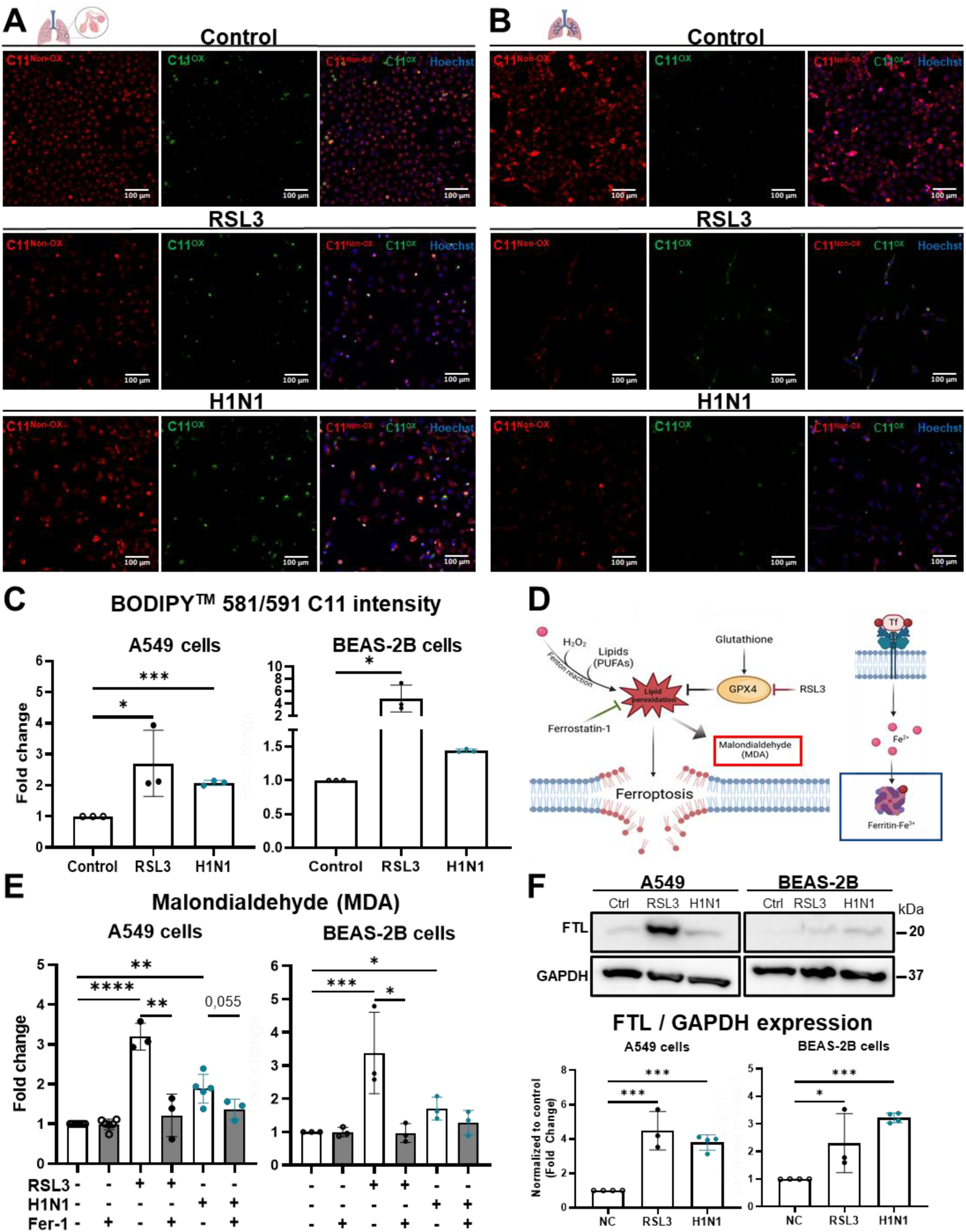
Influenza virus H1N1 causes increased levels of lipid peroxidation and FTL expression in A549 and BEAS-2B cells. Fluorescence images taken at a nominal magnification of 20x of (A) A549 and (B) BEAS-2B cells treated with RSL3 (A549: 10 µM and BEAS-2B: 250 nM) or infected with H1N1/PR8 influenza virus at MOI 1 stained with BODIPY^TM^ 581/591 C11 at 24 hpi and counterstained with Hoechst 33258. Red: non-oxidized lipids (C11^Non-Ox^); Green: oxidized lipids (C11^Ox^) Blue: nucleus (scale bar = 100 µm). Quantitative analysis of mean integrated fluorescence intensity of C11^Ox^/C11^Non-Ox^ ratio was measured for both (C) A549 and BEAS-2B cells (mean ± SD). Each dot represents the quantification of one biological replicate. At least 10 images per condition per biological replicate were analyzed. C11^Ox^/C11^Non-Ox^ ratio of control was defined as 100%. (D) Schematic representation of iron participation in Fenton reaction leading to oxidation of polyunsaturated fatty acids (PUFAs), the production of malondialdehyde (MDA), and representation of iron transport into the cell by transferrin receptor (Tf), as well as the subsequent storage of intracellular iron in ferritin. (E) MDA levels of A549 and BEAS-2B cell lysates collected at 24 hpi with influenza H1N1/PR8 and treatment with RSL3 (A549:10 µM and BEAS-2B: 250 nM) and pre-treatment with Fer-1 (5 µM). MDA concentrations were normalized to the corresponding protein concentration and are presented as fold changes to the control. (F) Representative blot showing the relative expression of FTL in A549 and BEAS-2B cells infected with influenza H1N1/PR8 at a MOI = 1 or treated with RSL3 (A549: 30 µM and BEAS-2B: 300 nM). Cell lysates were harvested at 24 hpi for Western blot analysis. Statistical analysis was performed using one-way ANOVA followed by Dunnett’s multiple comparisons test. *p < 0.05, **p < 0.01 ***p < 0.001. Data are represented as mean ± SD of at least three independent biological replicates.

In A549 cells, H1N1 infection increased the ratio of green to red signal (2.1 ± 0.1 fold) compared to the control group. As expected, the RSL3 treatment caused a more pronounced increase (2.7 ± 1.1 fold) in lipid peroxidation (Fig. 3C, left). Similarly, in BEAS-2B cells, H1N1 infection increased the ratio of green to red signal (1.4 ± 0.1 fold) compared to the control, while RSL3 treatment caused a marked increase (4.8 ± 2.2 fold) (Fig. 3C, right). To confirm whether the increased lipid peroxidation was a result of ferroptosis, the ferroptosis inhibitor Ferrostatin-1 (Fer-1) was used to pre-treat cells for one hour prior to infection or RSL3 treatment followed by the determination of malondialdehyde (MDA) as a measure of lipid peroxidation [53] (Fig. 3D). In A549 cells, MDA levels significantly increased in H1N1-infected (1.9 ± 0.4 fold) and RSL3-treated groups (3.2 ± 0.3 fold) compared to the control condition (Fig. 3E, left). In BEAS-2B cells, RSL3 treatment induced a similar increase in MDA levels (3.4 ± 1.2 fold) and H1N1 infection caused a lower but significant increase (1.7 ± 0.3 fold) (Fig. 3E, right). As expected, Fer-1 pre-treatment prevented the increase of MDA induced by RSL3 in both cell lines (Fig. 3E). Infected BEAS-2B and A549 cells pre-treated with Fer-1 also showed a lower level of MDA with 1.3 ± 0.4 fold and 1.4 ± 0.3 fold compared to untreated infected cells (1.7 ± 0.3 and 1.9 ± 0.4 fold), respectively. However, this difference did not reach statistical significance.

The excess oxidation of PUFAs is driven by labile, ferrous iron (Fe^2+^) participating in the Fenton reaction, indicating that processes involved in the import, export, and storage of iron must be tightly regulated to prevent lipid peroxidation [51]. Ferritin, the major iron storage protein in cells (Fig. 3D), is known to play an important role in ferroptosis and in the replication of H1N1 [19], therefore, we next determined whether ferritin expression levels were altered in H1N1-infected cells. In A549, ferritin expression levels significantly increased (3.8 ± 0.4 fold) in H1N1-infected cells (Fig. 3F), a change comparable to that in the RSL3-treated group (4.5 ± 1.1 fold) relative to the control. Similarly, in BEAS-2B cells infected with H1N1, ferritin expression levels significantly increased (3.2 ± 0.2 fold) compared to the control. This increase in ferritin expression levels of infected cells confirms a disruption of intracellular iron homeostasis and suggests a compensating mechanism as a response to an increase in intracellular labile iron. Consistent with previous reports [17, 19], our results indicate H1N1 influenza virus infection causes ferroptosis features, as evidenced by increased lipid peroxidation, MDA levels and ferritin protein expression in A549 and BEAS-2B cells.

### 3.2 Human adenovirus type C5 causes cell death at longer times of incubation

Next, we investigated whether ferroptosis features are also triggered by human adenovirus type C5 (HAdV-C5). To assess infection dynamics and to capture potential early infection events, cells were infected with HAdV-C5 at an MOI of 1 and the expression of hexon, a major structural protein found in the capsid of HAdV-C5, was monitored as a marker of infection. Since HAdV structural proteins can take over 36 hpi to become detectable [54], we stained for hexon protein at 24, 48, 72 and 96 hpi (supplementary Fig. 2).

In A549 cells, it took over 48 hpi for most cells to become hexon^+^, with nearly all cells (93.48% ± 4.75%) being hexon^+^ by 72 hpi, whereas at 96 hpi only a few nuclei were visible, indicating cell lysis (Fig. 4A). In contrast, in BEAS-2B cells, it took over 72 hpi for the majority of cells (96.81% ± 4.36%) to become hexon^+^, indicating a slower infection rate in bronchial epithelial cells (Fig. 4B).

**Fig. 4.**
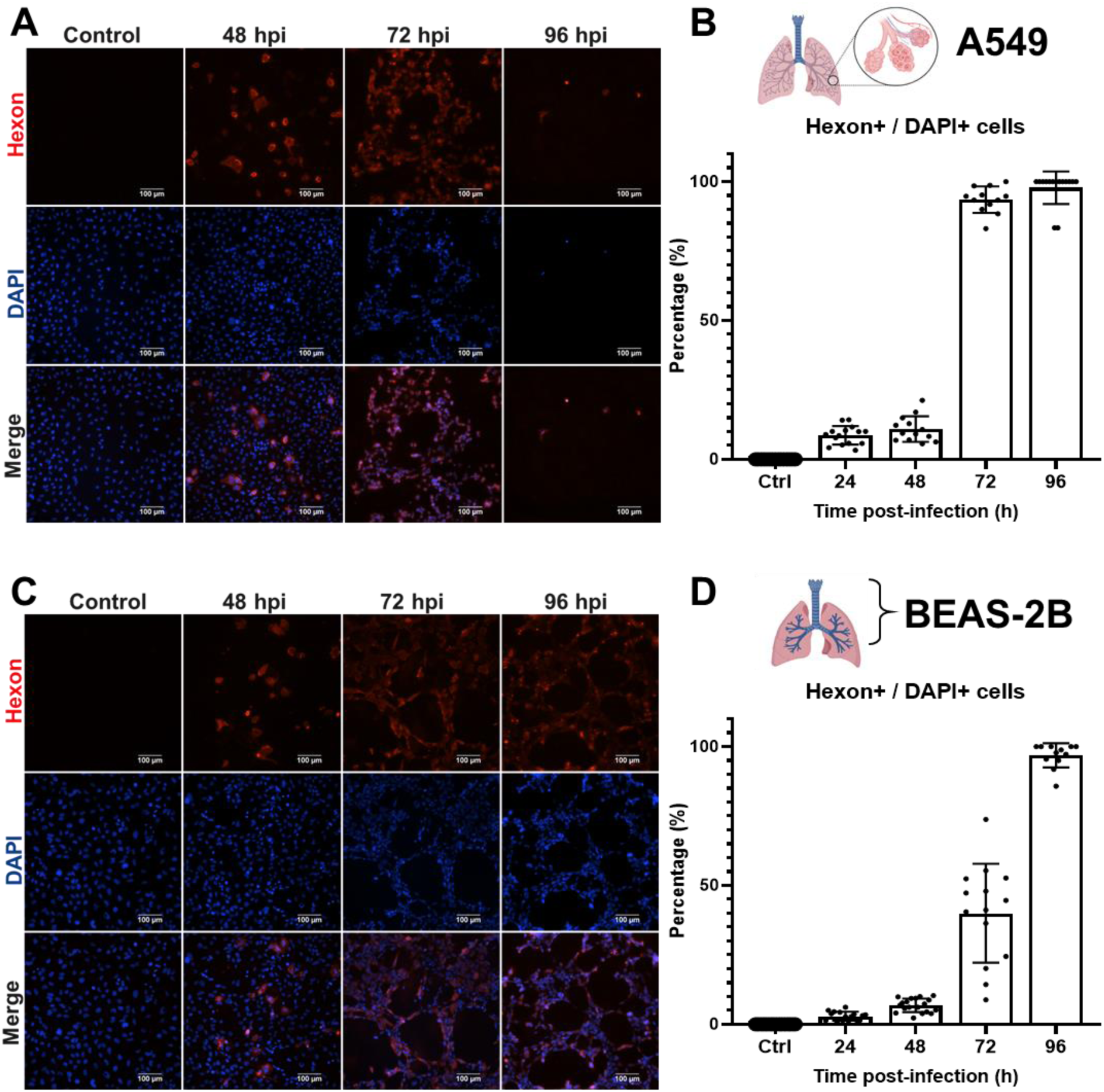
HAdV-C5 replication rate in lung epithelial cells. Fluorescence images taken at a nominal magnification of 20x of (A) A549 and (C) BEAS-2B cells infected with HAdV-C5 virus at a MOI = 1 fixed and stained at 48, 72 and 96 hpi with anti-hexon antibody and counterstained with DAPI. Red: hexon positive cells; Blue: nucleus (scale bar = 100 µm). Quantitative analysis of hexon positive and DAPI positive (B) A549 and (D) BEAS-2B cells was done on at least 10 images per condition using ImageJ. Results are shown as percentage of Hexon+/DAPI+ cells (mean ± SD).

At 48 hpi, morphological changes consistent with the cytopathic effect of HAdVs such as cell rounding and swelling, were observed in A549 (Fig. 5A, left). In contrast, comparable morphological changes were observed in BEAS-2B cells only at 72 hpi (Fig. 5A, right). PI staining revealed an increasing number of PI^+^ cells over time in A549, but not in BEAS-2B cells (Fig. 5B). Consistent with these findings, cell metabolic activity in A549 cells was significantly reduced (59.6% ± 6.1%) by 72 hpi (Fig. 5C). Quantification of PI^+^ cells showed a significant increase in cell death at 48 hpi (23.30% ± 7.83%), with a further increase at 72 hpi (40.77% ± 13.94%) (Fig. 5D). In BEAS-2B cells, metabolic activity remained unaffected up to 72 hpi, and PI quantification showed only a small increase in cell death at 72 hpi (18.0% ± 1.7%) (Fig. 5E and 5F).

**Fig. 5.**
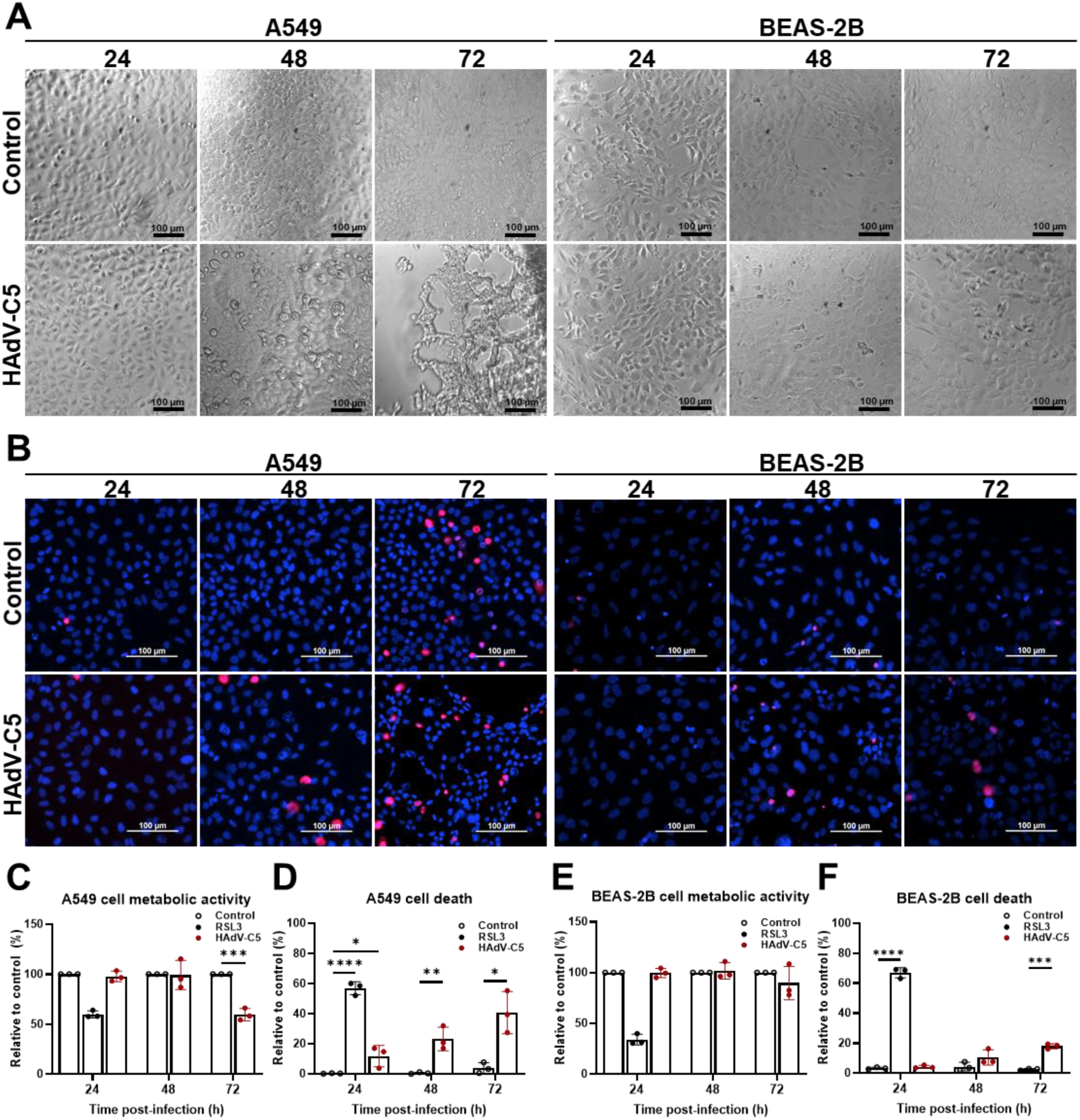
HAdV-C5-induced cell death has an earlier onset in A549 cells compared to BEAS-2B cells. (A) Representative brightfield microscope images of A549 and BEAS-2B cells infected with HAdV-C5 virus at a MOI = 1 at 24, 48, and 72 hpi (scale bar = 100 µm). (B) Representative fluorescence images of PI and Hoechst double staining taken at a nominal magnification of 20x. Red: PI positive, dead cells; Blue: nucleus (Scale bar = 100 µm). MTT assay as measurement of cell metabolic activity of viable cells of (C) A549 and (E) BEAS-2B cells. Cell metabolic activity of the control group was defined as 100% activity. Each dot represents a biological replicate with at least 6 technical replicates per condition. Results are shown as percentage relative to the control (mean ± SD). Cell death of (D) A549 and (F) BEAS-2B as measured by PI staining. RSL3 at 24 h was introduced as a single time point positive control. The number of dead cells was quantified according to the equation stated in the methods section. In short, the total number of dead cells is calculated as the sum of PI+ cells and the difference between the average number of DAPI+ cells in the control and the number of DAPI+ cells in the treatment, divided by the average number of DAPI+ cells in the control. Each dot represents the average of ten images per condition (mean ± SD). Statistical analysis was performed using one-way ANOVA followed by Dunnett’s multiple comparisons test. *p < 0.05, **p < 0.01, ***p < 0.001, ****p < 0.0001, ns: non-significant. Results are of at least three biological replicates.

Next, lipid peroxidation was measured by staining with BODIPY^TM^ 581/591 C11 following infection with HAdV-C5 at 24, 48 and 72 hpi in A549 (Fig. 6A) and BEAS-2B (Fig. 6B) cells. RSL3 treatment at 24 h served as a positive control (same as for H1N1 as the RSL3 and HAdV-C5 samples were prepared in the same experiment). Quantification of green to red fluorescence intensity ratios showed a time-dependent increase of oxidized lipids in A549 cells, with a significant change at 72 hpi (1.9 ± 0.4 fold) compared to controls. In contrast, no change in the ratio of oxidized to non-oxidized lipids was observed in BEAS-2B cells at any of the time points measured (Fig. 6C).

**Fig. 6.**
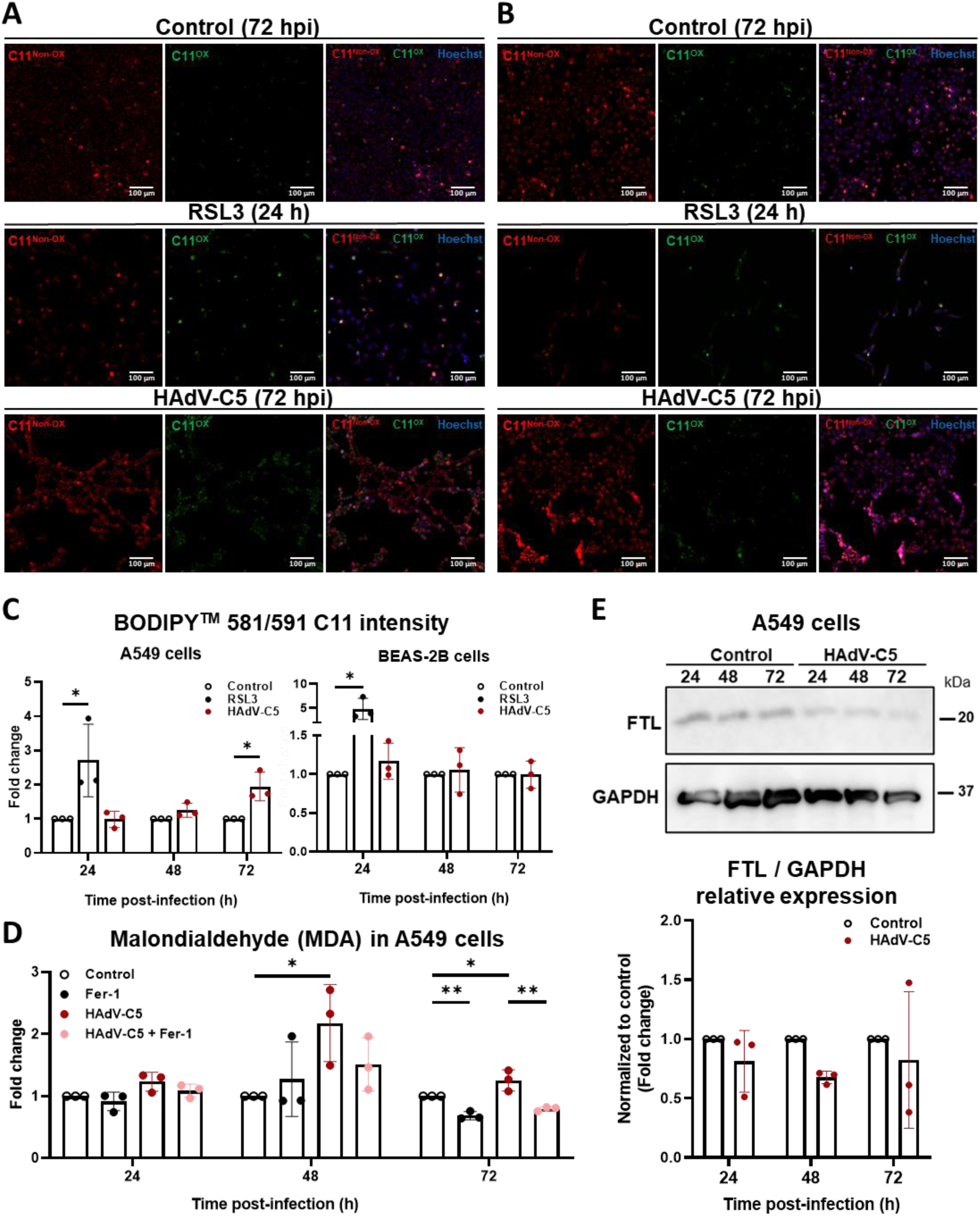
HAdV-C5-induced lipid peroxidation in A549 cells is time-dependent manner. Representative fluorescence images taken at a nominal magnification of 20x of (A) A549 and (B) BEAS-2B cells treated with RSL3 (A549:10 µM and BEAS-2B: 250 nM) for 24 h or infected with HAdV-C5 at a MOI = 1 stained with BODIPY^TM^ 581/591 C11 at 72 hpi and counterstained with Hoechst 33258. Red: non-oxidized lipids (C11^Non-Ox^); Green: oxidized lipids (C11^Ox^) Blue: nucleus (scale bar = 100 µm). (C) Quantitative analysis of mean integrated fluorescence intensity of C11^Ox^/C11^Non-Ox^ ratio was measured for both A549 and BEAS-2B cells (mean ± SD) at 24, 48 and 72 hpi. Each dot represents the quantification of one biological replicate. At least 10 images per condition per biological replicate were analyzed. C11^Ox^/C11^Non-Ox^ ratio of control was defined as 100%. Statistical analysis was performed using one-way ANOVA followed by Dunnett’s comparisons test. (D) MDA levels of A549 cell lysates collected at 24, 48 and 72 hpi with HAdV-C5 and treatment with RSL3 (10 µM) and pre-treatment with Fer-1 (5 µM). MDA concentration was normalized to the corresponding protein concentration and are presented as fold changes compared to the control. Data are represented as mean ± SD of at least three independent biological experiments with three technical replicates. Statistical analysis was performed using one-way ANOVA followed by Tukey’s post hoc multiple comparisons test. *p < 0.05, **p < 0.01.

Based on these findings, which indicate a higher susceptibility of A549 cells to HAdV-C5 infection, and the absence of observable changes in cell viability or lipid peroxidation in BEAS-2B cells, follow-up experiments were performed only on A549 cells. To further confirm the increase in lipid peroxidation, MDA levels were measured and assessed for sensitivity to the ferroptosis inhibitor Fer-1. MDA levels were significantly increased as early as 48 hpi (2.2 ± 0.6 fold) and remained significantly higher at 72 hpi (1.3 ± 0.2 fold) compared to the controls (Fig. 6D). Pre-treatment with Fer-1 significantly reduced the amount of MDA compared to non-Fer-1 pre-treated samples at 72 hpi in both control (0.7 ± 0.06 fold vs 1) and HAdV-C5 infected (0.8 ± 0.03 vs. 1.3 ± 0.2 fold) cells [19, 36]. Lastly, contrary to H1N1 infections of A549, no changes in the expression of FTL were found at any of the respective time points (Fig. 6E).

Overall, our data indicate that HAdV-C5 induces a marked increase in lipid peroxidation at later times of infection, which correlate with observable changes in cell morphology and hexon expression. However, the increase of MDA was only partially reduced by Fer-1, and no changes in FTL expression was detected, suggesting an iron-independent increase in lipid peroxidation.

### 3.3 H1N1 and HAdV-C5 infections cause alterations to the mitochondrial network in bronchial epithelial cells

Mitochondria regulate lipid metabolism, iron transport, and reactive oxygen species (ROS) production and thus play an active role in triggering ferroptotic cell death [10]. When exposed to ferroptosis inducers, such as RSL3, the mitochondrial outer membrane ruptures, causing the mitochondria to appear compacted and fragmented [4, 10]. Viruses are known to manipulate mitochondrial dynamics and morphology to benefit their own replication and survival [55, 56]. Therefore, we aimed to determine the effect of H1N1 and HAdV-C5 infection on the mitochondrial network and changes in morphology.

Morphological changes to mitochondria have been observed in H1N1 infected A549 cells at 24 hpi [18, 57]. To recapitulate this phenotype, we stained cells with MitoTracker^TM^ Red FM and visualized the mitochondrial network with fluorescence microscopy images, which were binarized using the *Mitochondria Analyzer* plugin [43, 48] for the quantification of mitochondria morphology and the *MiNA* plugin [43, 47] for the quantification of the footprint (area) occupied by mitochondria. We observed that mitochondria formed long, tubular-like structures in control A549 cells (Fig. 7A). In comparison, mitochondria in the RSL3-treated groups appeared fragmented and dot-like (Fig. 7A), observations confirmed by the significant decrease in mitochondrial footprint (14.4 ± 8.9 µm^2^ vs. 24.8 ± 6.8 µm^2^), branch length per mitochondrion (0.76 ± 0.2 µm vs.1.2 ± 0.2 µm), number of branches per mitochondrion (1.2 a.u. ± 0.1 vs. 1.3 ± 0.1 a.u), junctions per mitochondrion (0.07 ± 0.04 a.u.vs. 0.1 ± 0.06 a.u.) and mean form factor (1.5 ± 0.1 a.u. vs.1.8 ± 0.1 a.u.) indicating less and smaller, rounder mitochondria (Fig. 7B). In contrast, only small changes of mitochondrial network structure could be seen in H1N1 and HAdV-C5-infected A549 cells with more doughnut-shaped mitochondria appearing compared to the control. However, quantification confirmed no significant changes in mitochondrial footprint and morphology parameters, such as branch length and junctions per mitochondrion and mean form factor, caused by the viruses at 24 hpi (Fig. 7B).

**Fig. 7.**
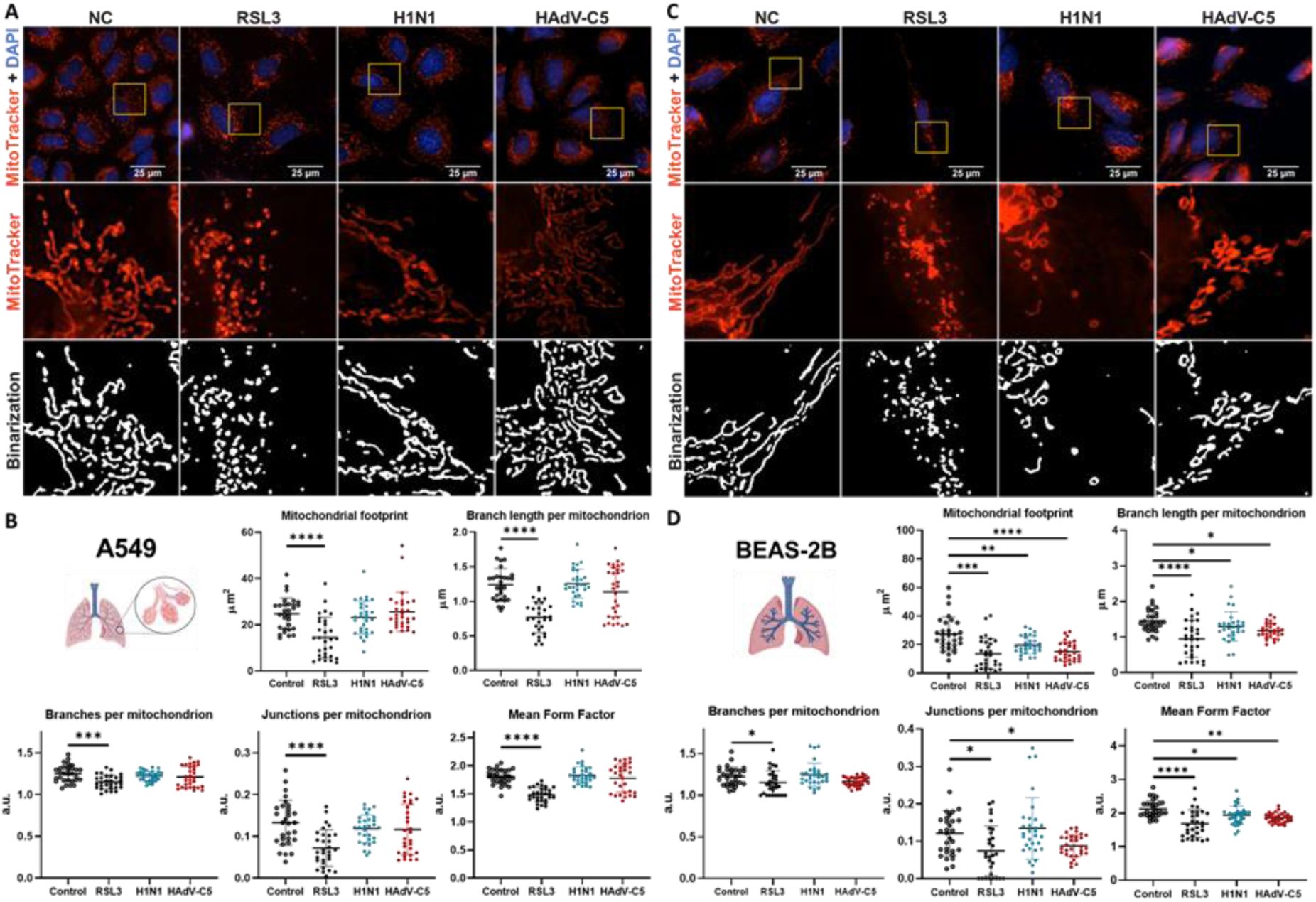
Influenza H1N1 and HAdV-C5 infections cause alterations in mitochondrial morphology in BEAS-2B but not A549 cells. MitoTracker^TM^ Red FM fluorescence images of (A) A549 and (C) BEAS-2B cells at a nominal magnification of 60X after treatment with RSL3 (A549: 10 µM; BEAS-2B: 250 nM) for 24 h or infection with influenza H1N1/PR8 or HAdV-C5 at a MOI = 1 at 24 hpi Red: mitochondria; Blue: nucleus (scale bar = 25 µm). Enlarged images of boxed area are shown for MitoTracker^TM^ and after binarization using the *Mitochondria Analyzer* plugin [43, 48]. Parameters of mitochondrial network analysis for (B) A549 and (D) BEAS-2B cells. Mitochondrial footprint of at least 10 single cells was analyzed for each condition (mean ± SD). Branch length per mitochondrion, branches per mitochondrion, junctions per mitochondrion and mean form factor of at least 10 whole images per condition was determined (mean ± SD). Statistical analysis was performed using one-way ANOVA followed by Dunnett’s multiple comparisons test. *p < 0.05, **p < 0.01, ***p < 0.001, ****p < 0.0001. Results are of at least three biological replicates.

Interestingly, BEAS-2B cells infected with H1N1 had mitochondria that appeared shorter and doughnut-shaped (Fig. 7C). Indeed, quantification of mitochondrial footprint and morphology parameters showed a significant decrease in mitochondrial footprint (20 ± 6 µm^2^ vs. 27 ± 12 µm^2^), branch length per mitochondrion (1.3 ± 0.4 µm vs. 1.5 ± 0.3 µm), and mean form factor (1.9 ± 0.3 a.u. vs. 2.1 ± 0.3 a.u.) indicating rounder mitochondria with shorter branches compared to controls (Fig. 7D). Similarly, BEAS-2B cells infected with HAdV-C5 exhibited shorter mitochondrial branches (Fig. 7C), which was reflected by the significant reduction in mitochondrial footprint (15 ± 7 µm^2^ vs. 27 ± 12 µm^2^), branch length per mitochondrion (1.2 ± 0.2 µm vs. 1.5 ± 0.3 µm), junctions per mitochondrion (0.09 ± 0.03 a.u. vs. 0.12 ± 0.06 a.u.) and mean form factor (1.9 ± 0.1 a.u. vs. 2.1 ± 0.3 a.u) compared to controls (Fig. 7D). Profound changes could be seen in the RSL3 treated cells which appeared fragmented and dot-like. These observations were confirmed by the significant reduction in mitochondrial footprint (13.6 ± 10.7 µm^2^ vs. 27 ± 12 µm^2^), branch length per mitochondrion (0.95 ± 0.5 µm vs. 1.5 ± 0.3 µm), branches per mitochondrion (1.2 ± 0.1 a.u. vs. 1.2 ± 0.1 a.u.), junctions per mitochondrion (0.08 ± 0.07 a.u. vs. 0.12 ± 0.06 a.u.) and mean form factor (1.7 ± 0.4 a.u. vs. 2.1 ± 0.3 a.u) compared to controls (Fig. 7D)

Since the changes caused by HAdV-C5 infection were observed only at later times of infection, we stained the mitochondria of A549 infected cells at 48, 60 and 72 hpi. Images showed a clear reduction of tubular-like structures and an increase in fragmented, dot-like mitochondria that could be observed in the HAdV-C5 infected cells at 48 hpi compared to its control group. By 72 hpi, only a few mitochondria could be observed in the infected cells, all of which appeared dot-like and compacted compared to the control cells at the same time point. The progressive deterioration in the mitochondria morphology (supplementary Fig. 3), coincided with time points in which we observed the increase in lipid peroxidation and cell death. However, accurate quantification of mitochondrial footprint and morphology parameters was not possible due to the mitochondria signal spanning over multiple imaging layers [48].

Overall, our results indicate that H1N1 and HAdV-C5 infection decrease mitochondrial footprint in BEAS-2B cells and cause detrimental changes to mitochondrial morphology. The level of mitochondrial alterations in these cells was comparable to the damage caused by RSL3.

## 4. Discussion

In this study we investigated ferroptosis-like hallmarks induced by H1N1 and HAdV-C5 infections in BEAS-2B (bronchial) and A549 (alveolar) epithelial cells. We provide evidence that influenza A virus (H1N1) induces lipid peroxidation and disrupts iron metabolism (Fig. 3), two key hallmarks of ferroptosis. Our findings are in line with earlier reports [17, 19]. Further studies reported a downregulation of ferritin heavy chain (FTH1) expression in A549 cells infected with H1N1/PR8 at a low MOI (0.1) [19], whereas our data shows an increase in the expression of ferritin light chain (FTL) in both A549 and BEAS-2B cells following infection with H1N1/PR8 at a higher MOI 1. Proteomics data of normal human bronchial epithelial (NHBE) cells infected with different strains of IAV at an MOI 2 indicate that the relative expression levels of FTH1 fluctuate in a time-dependent manner [58]. Indeed, clinical studies showed, using direct quantification methods of serum ferritin level, that high ferritin levels are linked to a more severe infection and can be used as a marker for severe complications and poor patient outcomes [59, 60]. Furthermore, the upregulation in FTL was comparable to that seen in RSL3-treated groups, suggesting a potential compensatory response, likely resulting from the upregulated expression of transferrin and transferrin receptor that has been previously reported in H1N1-infected A549 cells [17].

We further demonstrate, for the first time, that HAdV-C5 infection also induces lipid peroxidation in A549 cells at later times of infection (48 and 72 hpi) but not at early time points of infection (24 hpi) (Fig. 6) as indicated by the quantification of hexon protein expression (Fig. 4). These findings indicate that lipid peroxidation is a late-stage event in HAdV-C5 infection, rather than an early response. Strikingly, we did not see an increase of lipid peroxidation in BEAS-2B cells within the time frame of our experiments. However, given HAdV-C5 replication kinetics and onset of cell death in BEAS-2B cells are different compared to A549 cells, it may indicate that cellular pathways may be regulated differently eliciting an RCD pathway independent of ferroptosis-like hallmarks. Another explanation is that HAdVs elicit different innate and inflammatory responses during infection in different airway epithelial cells [61, 62], some of which may suppress lipid peroxidation cascades.

Interestingly, previous studies, performed in NHI3T3 fibroblasts, have shown that expression of the adenovirus early region 1A (E1A) gene can block the expression of ferritin in response to oxidative stress [29], further showing differential regulation of ferroptosis-like hallmarks in different cell types. We observed a trend towards a decrease in the expression of FTL in A549 cells infected with HAdV-C5. Whether this trend reflects E1A-mediated suppression of ferritin or the increase in lipid peroxidation observed is unrelated to iron metabolism remains to be clarified.

Beyond lipid peroxidation and ferritin expression, we also explored virus-induced alterations in mitochondrial footprint and morphology. Mitochondria actively contribute to the onset of cysteine-deprivation-induced ferroptosis through glutathione depletion and consequently the generation of lipid peroxides [11]. Erastin-induced ferroptosis dramatically changes mitochondria morphology, causing an increase of mtROS, leading to loss of mitochondrial membrane potential [63]. The influenza A virus PB1-F2 protein is known to induce mitophagy by translocating to mitochondria and mediating its degradation by binding to LC-3 to promote degradation and elimination of mitochondrial antiviral signaling proteins (MAVS) to impair innate immunity [64–66]. Here, we show a decrease in mitochondrial content and morphology in BEAS-2B cells but not in A549 cells, following H1N1 infection at 24 hpi (Fig.7). This cell-type-specific effect may be caused by the distinct pattern of sialic acids on bronchial epithelial cells, which promote H1N1 uptake, [14, 67] and therefore faster infection in BEAS-2B cells compared to A549 cells (Fig. 1). Furthermore, a study on mitochondrial morphology changes in H1N1 infected A549 cells also found distinct fragmentation patterns under starvation conditions, confirming the findings presented in the present study [57].

Adenoviruses are also known to modulate mitochondrial homeostasis to enhance virus replication [68–70]. The viral E1B-19k protein localizes to mitochondria, where it binds to the pro-apoptotic proteins Bax and Bak to prevent apoptosis [69, 71–74]. Recently, the viral minor capsid protein VI was shown to localize to mitochondria during late stages of infection and promote, via its membrane-lytic function, the release of mitochondrial DNA (mtDNA) and heat shock protein 60 (HSP60) into the cytosol [75], both of which are important for maintaining mitochondrial integrity [76]. We observed a decrease in mitochondrial footprint and changes in mitochondrial morphology in BEAS-2B cells infected with HAdV-C5 at early times of infection (24 hpi) that could not be seen in A549 cells at the same time point (Fig. 7). Further suggesting that adenovirus-induced disruption of mitochondrial homeostasis is both cell type- and time-dependent and is likely also connected to the lytic function of protein VI [75]. Indeed, the drastic changes in mitochondrial morphology at later time points post HAdV-C5 infection prevented reliable quantification of changes in mitochondrial network, as diffusion of the MitoTracker signal did not allow for accurate determination of the parameters, which is a known limitation of the software as specified by the developer [48].

Recent research shows that mitochondrial integrity is regulated by the localization of cyclic GMP-AMP (cGAMP) synthase (cGAS) to the outer mitochondrial membrane where it associates with dynamin-related protein 1 (DRP1), one of the known key regulators of mitochondrial fission, and facilitates its oligomerization to control excessive ROS production and limit lipid peroxidation, thereby inhibiting ferroptosis [77]. Importantly, cGAS, a cytosolic DNA sensor that following its binding to DNA catalyzes the synthesis of second messenger cGAMP to activate stimulator of interferon genes (STING) and the downstream production of interferons [78], was found to be significantly increased in cancer cells, including A549 cells [77]. The higher cGAS expression in A549 cells may explain why we only observed smaller changes in the mitochondrial network in H1N1- and HAdV-C5-infected cells at 24 hpi. In addition, levels of DRP1 were shown to be differentially altered during IAV infection at early and late stages of infection [57, 79]. Interestingly, it was further shown that increasing cyclic adenosine monophosphate (cAMP) levels activate protein kinase A (PKA), which is subsequently translocated to the outer mitochondrial membrane via A-kinase anchoring protein (AKAP), where it can phosphorylate DRP1 thereby inhibiting its function [80]. Importantly, it was previously shown that mitochondrial integrity and function is linked to cAMP levels and signaling in lung epithelial cells [35, 81]. Indeed, other studies showed that inhibition of either exchange protein activated by cAMP 1 (EPAC1) or phosphodiesterase 4 (PDE4) reduces IAV replication efficacy, suggesting a potential link between cAMP levels [82, 83], mitochondrial function and morphology, and viral replication. Strikingly, it was shown that HAdV infections can hijack cAMP signaling via E1A-mediated mimicry of AKAP functionality and binding of PKA, suggesting a similar mechanism as for IAV [84]. However, our knowledge on HAdV induced changes in DRP1 or other mitochondrial integrity regulators is limited. Therefore, future studies should address how viruses alter cAMP-linked signaling and its impact on mitochondrial function.

## 5. Conclusions

In conclusion, we confirm previous findings regarding the role of ferroptosis during H1N1 infection by demonstrating increased lipid peroxidation, elevated MDA levels, and altered FTL expression in two different lung epithelial cell types. Importantly, we provide novel evidence that HAdV-C5 infection induces lipid peroxidation and MDA accumulation at later stages of infection, a process that is prevented by pre-treatment with the ferroptosis inhibitor Fer-1 only at advanced stages of infection and is not accompanied by significant changes in FTL expression. Additionally, we show for the first time that both H1N1 and HAdV-C5 infections lead to significant changes in mitochondrial content and morphology in bronchial (BEAS-2B) epithelial cells at early times of infection comparable to mitochondrial changes caused by RSL3. Contrastingly, in alveolar A549 cells significant mitochondrial changes were observed only at later times of infection with HAdV-C5, whereas H1N1 induced changes recapitulate phenotypes that were seen before [57]. Taken together our comparative study of H1N1 and HAdV-C5 induced ferroptosis-like hallmarks, offer a more extended and detailed understanding of how lipid peroxidation, iron homeostasis, and mitochondrial morphology are altered. However, whether the cell death caused by them is ferroptosis, or a blend of various regulated cell death pathways will require further investigation.

## Abbreviations

AKAP: A-kinase anchoring protein
cAMP: cyclic AMP
cGAS: cyclic GMP-AMP synthase
cGAMP: cyclic GMP-AMP
DMSO: dimethyl sulfoxide
DRP1: dynamin-related protein 1
EPAC: exchange protein activated by
cAMP 1 E1A: adenovirus early region 1A
FTH1: ferritin heavy chain
FTL: ferritin light chain
GPX4: glutathione peroxidase 4
GSH: glutathione
HAdV: human adenovirus
HAdV-C5: human adenovirus type C5
hpi: hours post infection
HSP60: heat shock protein 60
IAV: influenza A virus
MAVS: mitochondrial antiviral signaling proteins
MDA: malondialdehyde
MOI: multiplicity of infection
MTT: 3-(4,5-Dimethyl-2-thiazolyl)-2,5-diphenyl-2H-tetrazolium bromide
mtROS: mitochondrial ROS
mtDNA: mitocondrial DNA
NCOA4: nuclear receptor coactivator 4
NP: nucleoprotein
PBS: phosphate buffered saline
PDE4: phosphodiesterase 4
PFA: paraformaldehyde
PI: propidium iodide
PKA: protein kinase A
PUFA: polyunsaturated fatty acid
pVI: viral minor capsid protein VI
RCD: regulated cell death
RIPK1: receptor interacting protein kinase 1
ROS: reactive oxygen species
RT: room temperature
STING: stimulator of interferon genes
TBA: 2-thiobarbituric acid
TBARS: thiobarbituric acid reactive substances assay
TCA: trichloroacetic acid

## Acknowledgments

A.L.M.C is a recipient of a Secretaría de Ciencia, Humanidades, Tecnología e Innovación (SECIHTI) scholarship (2021-000022-01EXTF-00086). A.M.D. is the recipient of an Alzheimer Nederland grant (WE.03-2018-04, The Netherlands), Parkinson Fonds (The Netherlands), and a Rosalind Franklin Fellowship co-funded by the European Union and the University of Groningen. M.S. received support by Alzheimer Nederland grant WE.03-2019-05 and is supported by the Deutsche Forschungsgemeinschaft (IRTG1874DIAMICOM-SP2) and Novartis unrestricted grant 50199468. K.R. is supported by a ZonMW Off Road Grant (04510242410069). We thank Maria João Caiado and Prof. Wilfred den Dunnen for sharing the ferritin light chain antibody with us.

## Author contributions

A.L.M.C., C.H.T.J. M.v.d.V, A.T., and P. A., performed experiments. A. L. M. C. and J. W. analysed data. J. d. V. I. and A. H. supplied H1N1/PR8 and guided experimental set-up. A. L. M. C., A. M. D., M. S., and K. R. planned experiments and A. L. M. C., M. S., and K.R. conceptualized the story. A. L. M. C., M. S., and K.R. wrote the manuscript with input from all authors.

