## SupplementaryMaterial for "Respiratory viruses induce ferroptosis-like features in lung epithelial cells"

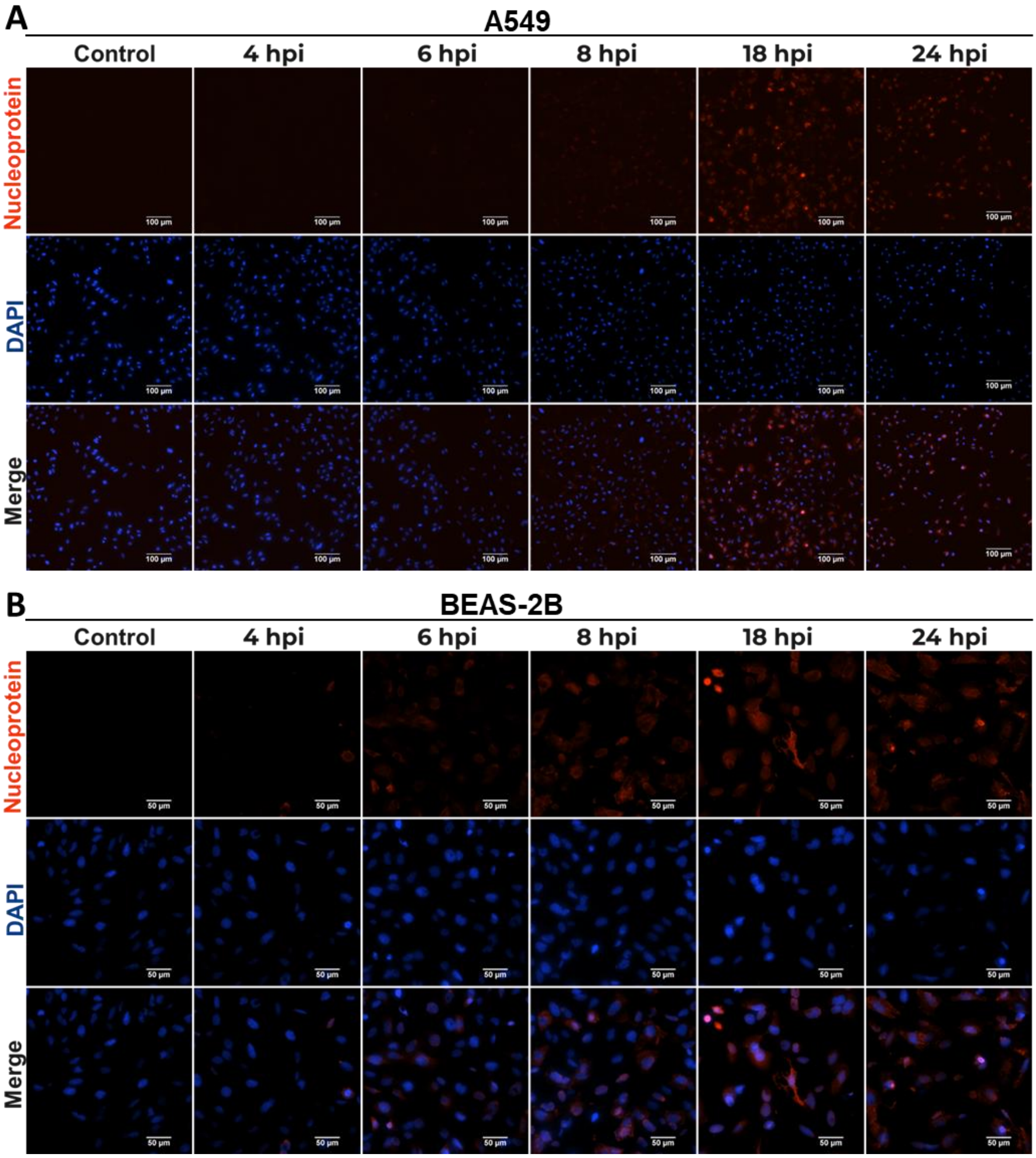


**Supplementary Fig. 1. H1N1 replication rate in lung epithelial cells**. Fluorescence images taken at a nominal magnification of 20x of (A) A549 and (B) BEAS-2B cells infected with H1N1/PR8 influenza virus at a MOI 1 fixed and stained at 4, 6, 8, 18 and 24 hpi with anti-influenza nucleoprotein (NP) antibody and counterstained with DAPI. Red: NP positive cells; Blue: nucleus (scale bar = 100 µm for A549 cells and 50 µm for BEAS-2B cells).

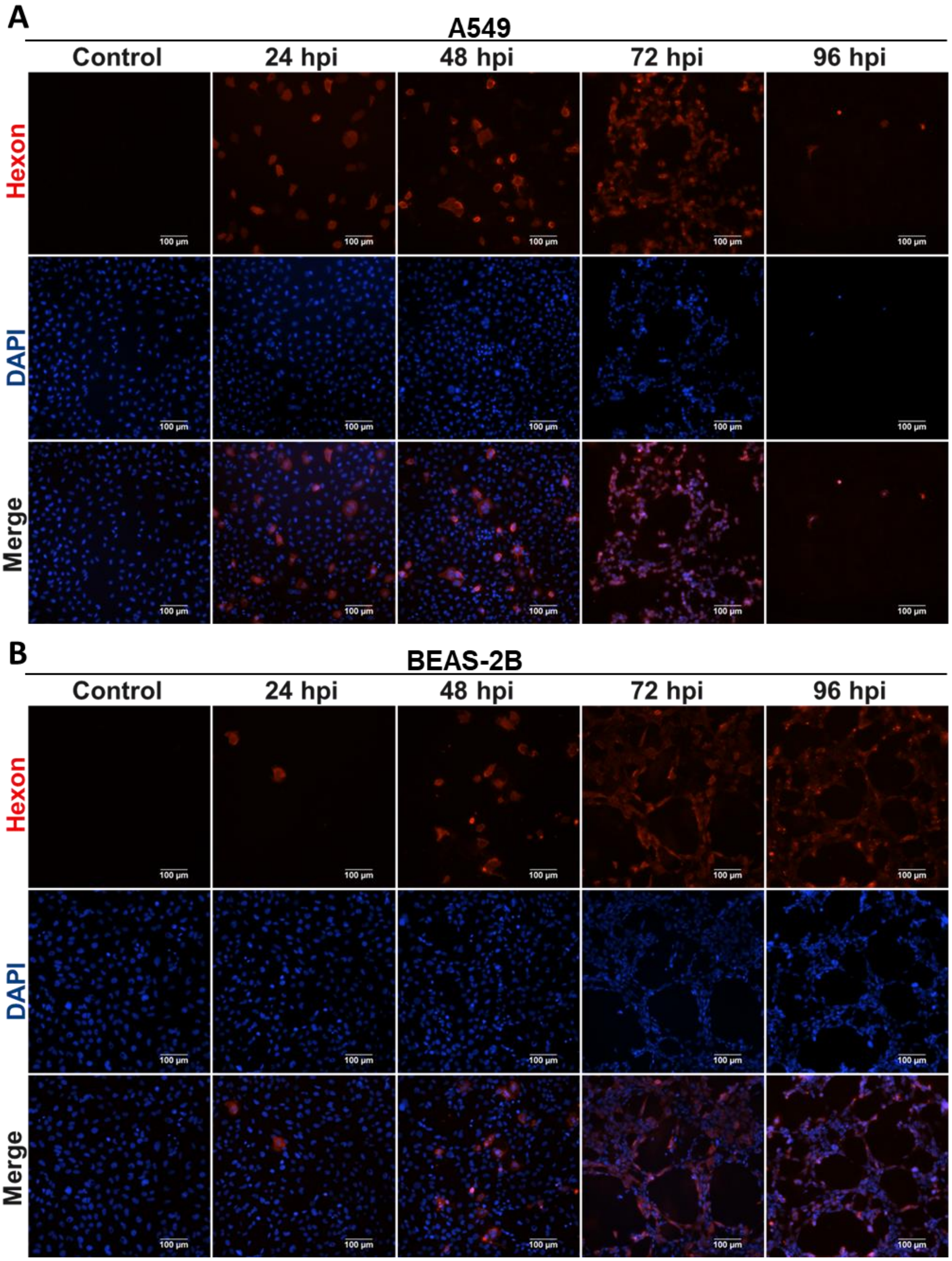


**Supplementary Fig. 2. HAdV-C5 replication rate in lung epithelial cells**. Fluorescence images taken at a nominal magnification of 20x of (A) A549 and BEAS-2B cells infected with HAdV-C5 virus at a MOI 1 fixed and stained at 24, 48, 72 and 96 hpi with anti-hexon antibody and counterstained with DAPI. Red: hexon positive cells; Blue: nucleus (scale bar = 100 µm).

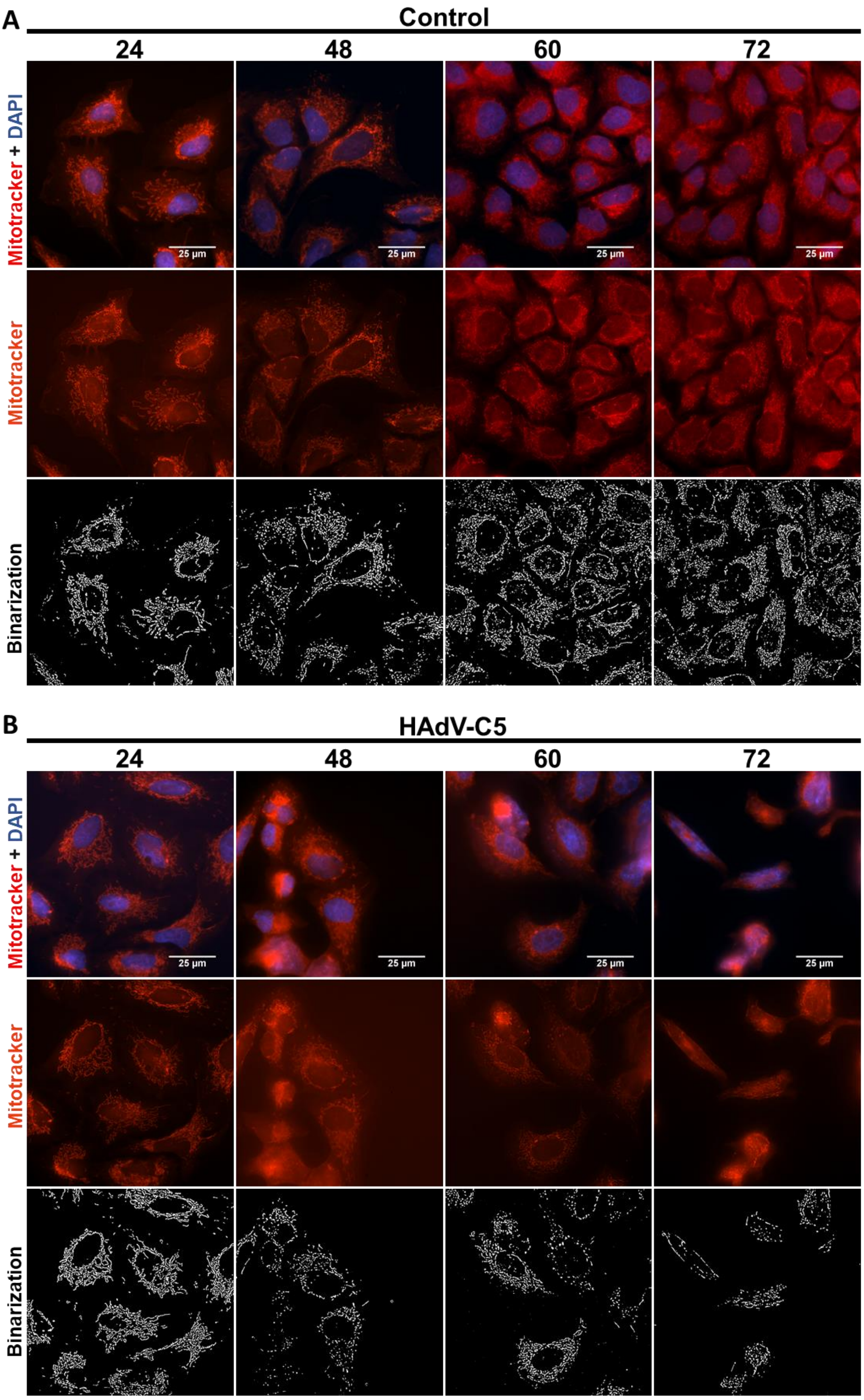
A
Control
24
48
60
72
Mitotracker + DAPI
Mitotracker
Binarization
25 µm
B
HAdV-C5
24
48
60
72
Mitotracker + DAPI
Mitotracker
Binarization
25 µm

**Supplementary Fig. 3. HAdV-C5 infection causes changes to mitochondria in A549 cells at later states of infection**. MitoTracker$^{TM}$ Red FM fluorescence images taken at a nominal magnification of 60X of A549 cells (A) control and (B) after infection with HAdV-C5 at an MOI 1 at 24, 48, 60 and 72 hpi Red: mitochondria; Blue: nucleus (scale bar = 25 μm).
